# TMPyP binding evokes a complex, tunable nanomechanical response in DNA

**DOI:** 10.1101/2023.10.16.562642

**Authors:** Balázs Kretzer, Levente Herényi, Gabriella Csík, Eszter Supala, Ádám Orosz, Hedvig Tordai, Bálint Kiss, Miklós Kellermayer

## Abstract

TMPyP is a porphyrin capable of DNA binding and used in photodynamic therapy and G-quadruplex stabilization. Despite its broad applications, the effect of TMPyP on DNA nanomechanics is unknown. Here we investigated, by manipulating λ-phage DNA with optical tweezers combined with microfluidics, how TMPyP influences DNA nanomechanics across a wide range of TMPyP concentration (5-5120 nM), mechanical force (0-100 pN), NaCl concentration (0.01-1 M) and pulling rate (0.2-20 μm/s). Complex responses were recorded, for the analysis of which we introduced a simple mathematical model. TMPyP binding leads to the lengthening and softening of dsDNA. dsDNA stability, measured as the force of DNA’s overstretch transition, increased at low (<10 nM) TMPyP concentrations, then decreased progressively upon increasing TMPyP concentration. The cooperativity of the overstretch transition decreased, due most likely to mechanical roadblocks of ssDNA-bound TMPyP. TMPyP binding increased ssDNA’s contour length. The addition of NaCl at high (1 M) concentration competed with many of the nanomechanical changes evoked by TMPyP. Because the largest amplitude of the nanomechanical changes are induced by TMPyP in the pharmacologically relevant nanomolar concentration range, this porphyrin derivative may be used to tune DNA’s structure and properties, hence control the myriad of biomolecular processes associated with DNA.

## Introduction

The advent and perfection of single-molecule methods in the recent decades have led to unprecedented new insights into the mechanisms and properties of biomolecular processes, and pointed out that mechanical forces and molecular nanomechanics play a much more important role in controlling cellular and sub-cellular phenomena than earlier thought. DNA has particularly been in the focus of single-molecule experiments, as the molecule lends itself naturally to exploring the role of its axial, bending, twisting and strand-separation nanomechanics in DNA-associated processes (1-5). The combination of novel methodologies, for example, optical tweezers, microfluidics and high-resolution fluorescence microscopies, has paved the way towards understanding how DNA nanomechanics are influenced by external factors such as intercalators (6), binding proteins (7) or solute concentration gradients (8). Understanding DNA nanomechanics is key not only to uncover the mechanisms of the vast array of DNA-related intracellular processes, for instance, DNA replication and repair, chromatin condensation, gene transcription and regulation, but also in the development of novel, high-efficacy pharmaceuticals that specifically and sensitively influence these DNA-dependent processes (9,10).

DNA nanomechanics are influenced by DNA-binding molecules that interact with DNA in a variety of ways (9-13). Certain proteins and small molecules bind to the major or minor grooves of dsDNA, thereby altering its structure (14) and stability (10,15-20). Others may display electrostatic or allosteric interactions with DNA, but neither of them results in the disrupting of genome continuity (10). Intercalators non-covalently insert their planar aromatic moieties between adjacent base pairs of dsDNA, thereby altering DNA’s structural and mechanical properties and perturbing enzymatic reactions crucial for cell proliferation and survival (6,10). Intercalation follows the rule of nearest-neighbor exclusion, hence adjacent base pairs are affected by every moiety intercalated (10,21). Intercalators typically lead to the stabilization, elongation and helix unwinding of dsDNA (10). Despite the pronounced structural and mechanical changes, intercalation is a reversible process, making intercalators promising candidates for a wide range of applications targeting DNA (6,10). By contrast, ssDNA-binding proteins and small molecules destabilize dsDNA (13,22,23). Altogether, DNA nanomechanics can be controlled by ligand binding *via* numerous synergistic and antagonistic mechanisms.

Porphyrins constitute an important and much investigated group of DNA-binding ligands, one of which, tetrakis(4-N-methyl)pyridyl-porphyrin (TMPyP), is the subject of the present paper. Porphyrins and their derivatives are widely known for their use in photodynamic tumor therapy (24-27). TMPyP, a cationic porphyrin, stands out by its strong affinity to DNA and by its fluorescent properties, making it potentially applicable in biosensing functions. Moreover, TMPyP and its derivatives are also used as building blocks in functional assemblies (28-31). Cationic porphyrin derivatives have a broad antimicrobial effect (24,32-35), and TMPyP has specifically been investigated for its virus-inactivating properties (36,37). The interaction of TMPyP with G-quadruplexes sparked major interest, as TMPyP alters the mechanical properties of the G-quadruplex, thereby perturbing telomerase activity (38-40), raising the serious possibility that TMPyP and its derivatives may be successfully employed in cancer treatment (24,41). Prior studies have shown that TMPyP can intercalate between base pairs due to its planar structure, and it can also bind to the minor groove of the dsDNA (36,37,42). The preferred type of binding depends on TMPyP concentration, ionic strength and DNA sequence; however, the simultaneous presence of the different binding modes can be detected even at low TMPyP/base pair ratios (36,37). TMPyP is also capable of binding to ssDNA, and it catalyzes the formation of dsDNA (43,44). The binding reactions occur on the 10-500 ms time scale, and intercalation has been shown to be slower than groove binding (45). Despite the extensive investigation of TMPyP binding to DNA, its effect on DNA nanomechanics is yet unknown.

In the present work we carried out a comprehensive analysis of the effects of a wide range of TMPyP concentration on the nanomechanical behavior, manifested in the force *versus* extension function, of DNA. Furthermore, we tested the effects of NaCl concentration and pulling rate on the TMPyP-dependent DNA nanomechanics. A complex array of nanomechanical response was measured, which we analyzed with a newly-developed empirical mathematical model that provided insight into the molecular mechanisms of the effects. Our results imply that DNA nanomechanics, hence the DNA-dependent processes, may be finely tuned by an interplay between nanomolar TMPyP concentrations and piconewton forces.

## Materials and Methods

### Samples and buffer solutions

For the entire set of nanomechanical measurements we used λ-phage dsDNA biotinylated at its 3’-3’ ends (Lumicks, Amsterdam, The Netherlands). To generate the data in **Figure 1**, λ-phage dsDNA biotinylated on its 5’-3’ ends was used (Lumicks, Amsterdam, The Netherlands). In all experiments, DNA was diluted to a final concentration of 20 ng/mL. DNA was tethered between two 3.11 μm diameter streptavidin-coated polystyrene microbeads (Kisker Biotech, Steinfurt, Germany). TMPyP (Porphychem, Dijon, France) was used at different concentrations indicated in the figures. Tris-HCl buffer (20 mmol/L Tris-HCl, pH 7.4) was used throughout the measurements. NaCl was added in different concentrations (0.01, 0.1 and 1 M). To inhibit the non-specific binding of positively charged TMPyP to the negatively charged glass surfaces, hence the alteration of the effective TMPyP concentration inside the flow cell, Tween-20 was added to the buffer at a final concentration of 0.01% (v/v). TMPyP concentration was measured by using a 4E UV-VIS absorption spectrophotometer (Varian, Inc., Palo Alto, California).

**Figure 1.**
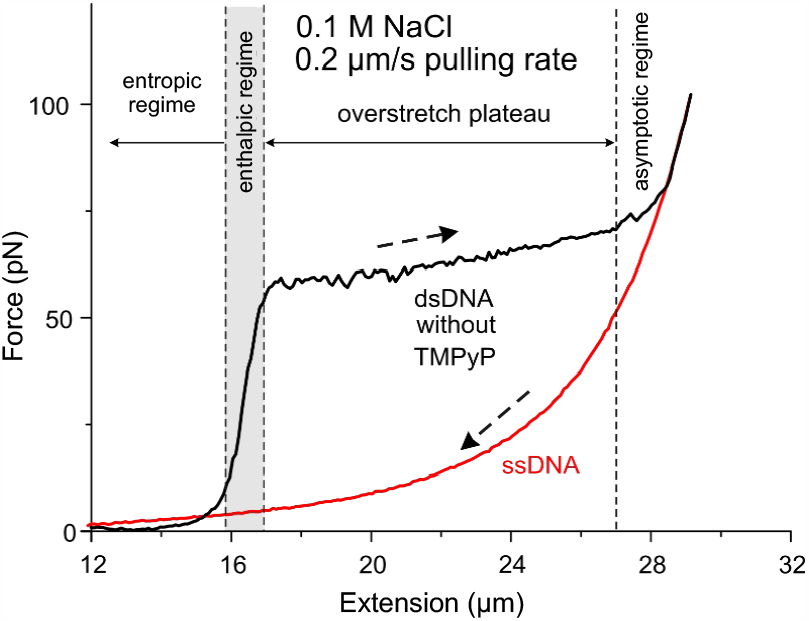
Canonical force *versus* extension curves (FECs) of dsDNA (black) and ssDNA (red). To obtain the FEC of dsDNA, one of the strands of a single λ-phage DNA molecule, was biotinylated on both the 3’ and 5’ ends and was extended with a constant rate (0.2 μm/s). The pulling was carried out in the absence of TMPyP, and in the presence of 0.1 M NaCl (20 mmol/L Tris-HCl, pH 7.4). At the end of the asymptotic part of the force trace we applied flow to wash off the complementary strand, and relaxed the remaining single-stranded DNA (ssDNA, red trace). Dashed arrows indicate the direction of data acquisition. The four characteristic nanomechanical regimes (entropic, enthalpic, overstretch plateau, asymptotic part) are separated by dashed vertical lines.

### Stretching single dsDNA molecules

Experiments were performed by using a dual-trap optical tweezers instrument coupled with a multi-channel microfluidic system (Lumicks, C-Trap, Amsterdam, The Netherlands). Streptavidin-coated microbeads were captured with the optical traps, and single molecules of biotinylated λ-phage dsDNA were tethered between them. By moving one of the beads, the tethered DNA was stretched and the resulting force was recorded. Following the simultaneous measurement of force and inter-bead distance, force-extension curves (FECs) of the single DNA molecules were plotted. To minimize TMPyP concentration fluctuations resulting from non-specific adsorption to the internal surfaces of the microfluidic chamber, the flow cell was incubated with the respective buffer for 45 minutes prior to the measurements. To obtain the force curve of a single DNA molecule, first it was pulled in buffer without TMPyP (control measurement) at a constant, pre-adjusted pulling rate. Subsequently, the molecule was brought to the microfluidic channel containing TMPyP at the given concentration, and it was positioned far from the channel wall so that diffusion-driven transport processes would not influence the effective TMPyP concentration (6,8). Then, the pulling cycle was repeated with the same rate. To alleviate hydrodynamic perturbations on DNA, flow was halted during the pulling cycle. Three sets of measurement were performed, each at different NaCl concentrations (0.01, 0.1 and 1 M). In a single measurement series we gradually increased the TMPyP concentration from 5 nM to 5120 nM, which resulted in ten measurement sets for each series. In each measurement set we collected several FECs at different pulling rates (0.2 μm/s, 2 μm/s, 20 μm/s). Thus, a multi-parametric dataset was recorded systematically with varying NaCl and TMPyP concentrations and pulling rates.

### AFM imaging of dsDNA

300-bp-long dsDNA fragments were imaged in liquid at 25 °C by using an Asylum Research Cypher ES atomic force microscope (Oxford Instruments, Abingdon, UK). Sample surfaces were scanned in tapping mode with BL-AC40TS (Olympus) cantilevers, resonated by photothermal excitation (∼20 kHz). Typical scan speeds were around 0.5 μm/s. Scanning resolution was 512 pixels/line for all images. 100 μl of DNA sample was dropped onto dried, poly-L-lysine (PLL) covered mica surface. To measure the effects of TMPyP, dsDNA previously bound to the PLL surface was incubated with 250 nM TMPyP for 10 min. Image post-processing and analysis were performed within the AFM driving software (IgorPro, WaveMetrics, Portland, OR, USA). The contour length of individual DNA strands (n=75) was measured for both control and TMPyP treated samples manually.

### Modeling the nanomechanical response of dsDNA

We developed an empirical mathematical model to fit the measured FECs and to understand which phases of dsDNA’s force response are most sensitive to a given experimental parameter. This model consists of three components: a sigmoid function (*f*_1_, entropic and enthalpic regimes), a linear function (*f*_2_, overstretch plateau) and a hyperbola (*f*_3_, asymptotic regime) (see **Figure 3a**):

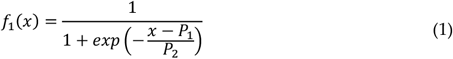

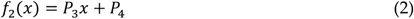

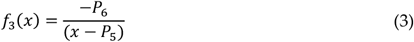

where the fitting parameters are marked with *P*_*i*_. The meaning of the individual parameters is the following:

□ *P*_1_: position of the sigmoid curve along the X-axis; scales with the contour length of dsDNA.
□ *P*_2_: step width of the sigmoid curve; scales with the compliance of dsDNA in the enthalpic region.
□ *P*_3_: slope of the linear function; scales inversely with the cooperativity of the overstretch transition.
□ *P*_4_: Y-axis intercept of the linear function; scales with the height of the overstretch plateau, hence with the stability of dsDNA.
□ *P*_5_: location of the asymptote of the hyperbola; scales with the maximal length of overstretched DNA (contour length of ssDNA).
□ *P*_6_: curvature of the hyperbola; related remotely to the bending rigidity of ssDNA.

Adding equations (2) and (3) and multiplying by equation (1) yields the following function:

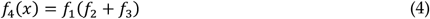

which was used to fit all the measured FECs.

### Data Analysis and Visualization

Raw data from the optical tweezers experiments were converted and analyzed using Python 3.9’s matplotlib 3.3.4 library and the lumicks.pylake 0.7.2 package. Fitting equation (4) to the data was carried out by using Origin (Northampton, Massachusetts, USA). Experimental force *versus* extension curves were fitted, by using IgorPro (version 9), with the non-extensible wormlike chain model,

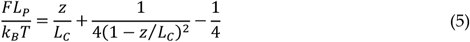

where *F, L*_P_, *k*_B_, *T, z* and *L*_C_ are force, persistence length, Boltzmann’s constant, absolute temperature, extension and contour length, respectively (46). Statistical data analysis of AFM images was done in R (version 4.1.0). Data visualization was carried out in Origin, CorelDraw, Inkscape (version 0.92), KaleidaGraph (version 5.0.5), IgorPro (version 9) and R (version 4.1.0).

## Results

### Reference force-extension curve of DNA

To assess the effects of TMPyP on the nanomechanical behavior of DNA, we first measured a reference force-extension curve (FEC) (**Figure 1**). This typical FEC reveals the nanomechanical behavior of double-stranded (ds) and single-stranded (ss) λ-DNA without any DNA-binding molecule present. In this experiment the dsDNA molecule was torsionally unconstrained and topologically open. We recorded a characteristic FEC that could be divided into four different regimes: i) entropic regime, ii) enthalpic regime, iii) overstretch plateau and iv) asymptotic regime. In the entropic regime the molecule is greatly extended by low (<10 pN) pulling forces which reduce configurational entropy, and the end-to-end distance of dsDNA approaches its contour length. In the enthalpic regime a linear force response is observed, the slope of which is related to the stretch modulus of dsDNA. In the overstretch plateau, dsDNA is extended beyond its native length at the expense of several structural transitions (B-S transition, melting-bubble formation, strand unpeeling) that occur cooperatively within a narrow, ionic strength-dependent force range (47). The features (height, length) of the overstretch plateau reflect the stability of dsDNA, hence they are sensitive to the presence of intercalators and groove-binding molecules (6,9-13,19,48-51), or ssDNA-binding proteins (13,22,23). In the final stage of the stretch force curve, the overstretch plateu is followed by an asymptotic regime, an elastic region where further elongation of DNA requires high forces. In this regime the two strands are held together by a few GC-rich regions (5). Since DNA was biotinylated on the 3’ and 5’ ends of the same strand, washing the mechanically dissociated DNA strand away and relaxing the molecular system yielded the FEC of the remaining ssDNA, which was characterized as a wormlike chain (**Figure 1**). The obtained reference FEC allows us to uncover the mechanistic details behind the effects of TMPyP binding to DNA.

### Force-extension curves in the TMPyP, NaCl, and pulling rate parameter space

We recorded dsDNA FECs across a wide range of TMPyP concentrations (doubling from 5 to 5120 nM) at three different NaCl concentrations (0.01, 0.1 and 1 M) and pulling rates (0.2, 2 and 20 μm/s), so that nine series of data were obtained (**Figure S1**). **Figure 2** highlights the essential features of the effects on the stretch force curves. Increasing TMPyP concentration lead to a drastic change in the FEC (**Figure 2.a**) so that all of the distinct regimes of the canonical FEC (**Figure 1**) were affected: the contour length of dsDNA increased, the slope in the enthalpic regime decreased, the height and slope of the overstretch transition decreased and increased, respectively, and the maximum contour length of the overstretched DNA increased. Upon increasing the pulling rate, we observed a slight recovery from the TMPyP effects (**Figure 2.b**), suggesting that some of the changes are influenced by the thermodynamics of the molecular system. Upon increasing the concentration of NaCl to 1 M, a significant recovery from the TMPyP effects was observed in the entropic and enthalpic regimes (**Figure 2.c**), indicating that ionic strength has a differential effect on the DNA-TMPyP interaction.

**Figure 2.**
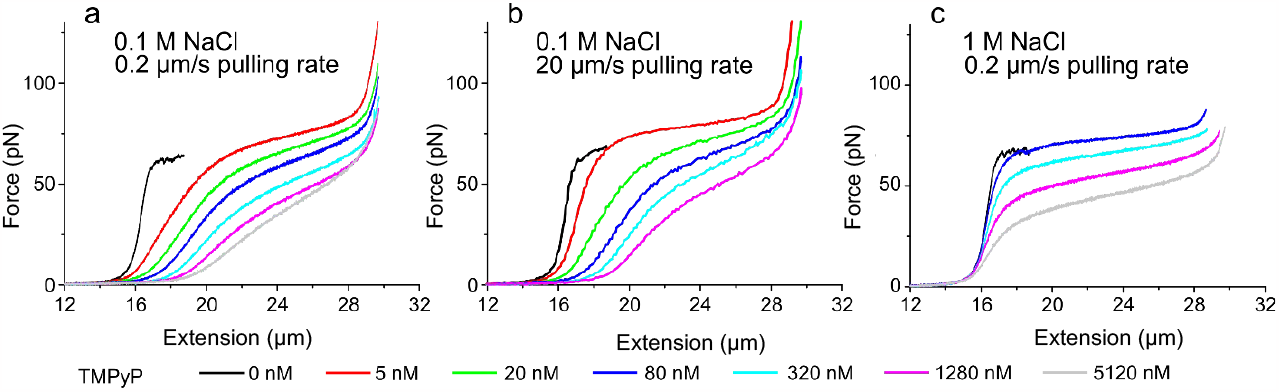
Force *versus* extension curves of λ-DNA stretched at different TMPyP concentrations (marked with different colors) in three different experimental conditions: **a)** 0.1 M NaCl and 0.2 μm/s pulling rate, **b)** 0.1 M NaCl and 20 μm/s pulling rate, **c)** 1 M NaCl and 0.2 μm/s pulling rate. The figure shows traces at a partial number of different TMPyP concentrations for clarity; for a complete FEC dataset obtained in this experimental parameter space, see **Figure S1**.

**Figure 3.**
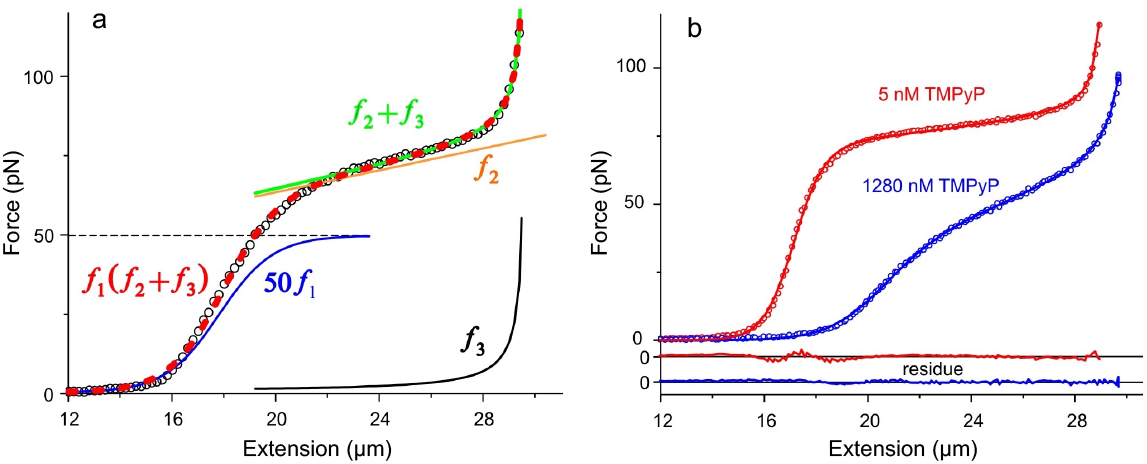
**a)** Schematics of applying the empirical mathematical model to the force *versus* extension curves. The measured FEC is marked with open black circles. The model comprises a sigmoid (blue curve; 50*f*_1_ magnified for better visualization), a linear function (yellow curve; *f*_2_), and a hyperbolic (black curve, *f*_3_), so that the fitting function is *f*_1_(*f*_2_ + *f*_3_) (red dashed curve). The equations of the functions are explained in the Results. *f*_2_ + *f*_3_ is shown with green continuous line. **b)** Illustration of the goodness of fit in two extreme cases of TMPyP concentration: 5nM (red) and blue: 1280nM (blue). Pulling was carried out at a NaCl concentration and pulling rate of 0.1 M and 20 μm/s, respectively. Open circles are the experimental data points, and the fits are marked with continuous lines. The colors of the residual traces correspond to those of the experimental data.

### Extraction of nanomechanical details with an empirical mathematical model

To dissect the effects of TMPyP, NaCl and pulling speed in detail, and to assign the effects to the structural and nanomechanical features of DNA, we developed an empirical mathematical model (equations 1-4) to fit the FECs with (**Figure 3**). The fitting function (equation 4) comprises three equations: a sigmoidal function (equation 1) that describes the transition from the entropic regime to the overstretch transition *via* the enthalpic regime; a linear function (equation 2) that describes the overstretch plateau itself; and a hyperbolic function (equation 3) that describes the asymptotic behavior of the FEC approaching maximal extension (**Figure 3.a**). The parameters (*P*_1-6_) of the equations scale with physical variables of DNA, as described in the Materials and Methods. **Figure 3.b** shows the fit of the model to the FECs measured at the extreme ends of the investigated TMPyP concentration range. The results show that the empirical model developed here fits the experimental data very well; therefore, plotting the fitting parameters as a function of TMPyP and NaCl concentrations and pulling rate is expected to reveal the mechanistic details of DNA’s nanomechanical response.

### Effect of TMPyP in the entropic regime

*P*1, which scales with the contour length of dsDNA, first increased rapidly, then slowly, as a function of increasing TMPyP concentration (**Figure 4.a**). The effect was completely alleviated by increasing the NaCl concentration to 1 M. Pulling rate had little effect on *P*1, detectable only at intermediate NaCl concentration (0.1 M) and at low (<40 nM) TMPyP concentrations (**Figure 4.a inset**). To investigate the TMPyP-induced DNA-lengthening effect in detail, we analyzed the length increment as a function of force **(Figures 4.b-c, S2**). The length increment increased monotonically as a function of force at every TMPyP concentration (**Figure 4.b**), suggesting that force enhances the binding of TMPyP to dsDNA. Extrapolating to zero force allowed us to calculate the TMPyP-induced DNA lengthening in the mechanically relaxed conformation. The DNA-lengthening effect of TMPyP could be substantiated with AFM measurements (**Figure S3**), even though its magnitude was smaller (9% *versus* 16%), due most likely to the constraints imposed by binding of DNA to the substrate surface. dsDNA length increased double-exponentially as a function of TMPyP concentration at forces between 0-45 pN (**Figure 4.c**). The apparent rate constants (k_1_, k_2_, see **Figure S2.b**) of the fitted double-exponential function differed by a factor of 20 and were force sensitive: the fast rate constant decreased, whereas the slow one increased linearly with force (**Figure S2.b**). The results imply that TMPyP binds by two different mechanisms with differing kinetic properties.

**Figure 4.**
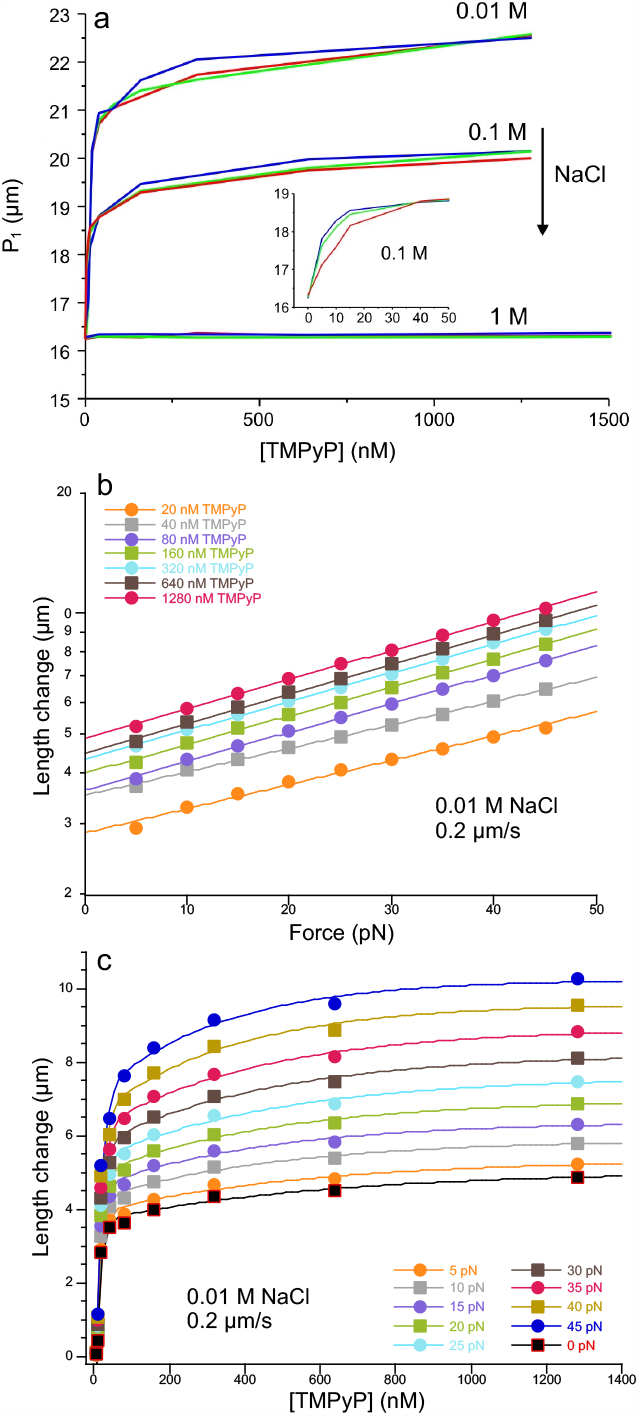
Effect of TMPyP on dsDNA nanomechanics in the entropic regime. **a)** P_1_ (sigmoid step position) as a function of TMPyP concentration, pulling rate and NaCl concentration (0.01, 0.1 and 1M). The pulling rates of 0.2 μm/s, 2 μm/s, 20 μm/s are indicated with blue, green and red, respectively. **Inset**, low-TMPyP-concentration (<50 nM) regime of the 0.1 M NaCl data. **b)** Length increment of dsDNA as a function of force at different TMPyP concentrations, in the presence of 0.01 M NaCl and at a pulling rate of 0.2 μm/s. The length change was calculated by subtracting the control (0 TMPyP) DNA length from the TMPyP-treated length measured at the given force. Data were fitted with single-exponential functions. **c)** Length increment of dsDNA as a function of TMPyP concentration at different forces, in the presence of 0.01 M NaCl and at a pulling rate of 0.2 μm/s. The zero-force data were obtained from the y-axis intercepts of **Figure 4.b**. Data were fitted with double-exponential functions. Analysis of the apparent rate constants of the fitting functions are shown in **Figure S2.b**.

### Effect of TMPyP in the enthalpic regime

*P*_2_, the width of the sigmoidal step, which scales with the apparent compliance of dsDNA, also increased rapidly, then slowly, as a function of increasing TMPyP concentration (**Figure 5.a**). The effect was significantly reduced, but it was not completely alleviated, by increasing NaCl concentration to 1 M. Pulling rate had little effect on *P*_2_, detectable only at intermediate NaCl concentration (0.1 M) and at low (<40 nM) TMPyP concentrations (**Figure 5.a inset**). Considering that in the enthalpic regime the dsDNA structure becomes distorted, we tested whether the molecular system is in equilibrium by comparing the stretch and relaxation force curves (**Figures 5.b-d**). At low NaCl concentration (0.01 M) and pulling rate (0.2 μm/s) we observed no force hysteresis across a wide TMPyP concentration range, indicating that the system was at thermodynamic equilibrium throughout the nanomechanical experiment (**Figure 5.b**). Upon increasing NaCl concentration to 1 M, a small hysteresis appeared at low TMPyP concentration (**Figure 5.c**). However, a systematic nanomechanical experiment, in which the maximum extension was progressively decreased, demonstrated that there is reversibility in the enthalpic regime, and hysteresis arises only if DNA has entered the overstretch transition (**Figure 5.d**). While *P*_2_ reflects the apparent compliance of dsDNA, the apparent stiffness is a more meaningful and accessible measure of the instantaneous nanomechanical behavior. To calculate the apparent, instantaneous longitudinal stiffness of the DNA molecule and the effect of TMPyP on this characteristic, we numerically derivated the FECs (**Figure 6**). In the absence of TMPyP a narrow bell-shaped curve, centered at 16.2 μm, was observed, independently of the NaCl concentration or the pulling rate. The peak position, hence the inflection point of the sigmoidal function (equation 1) coincides with the contour length of λ-phage DNA, substantiating the notion that the *P*_1_ parameter reflects the contour length of dsDNA in these experiments. The peak apparent stiffness of dsDNA is thus 50 pN/μm. Upon adding TMPyP at increasing concentrations, the curve broadened, the peak value decreased, and the peak position progressively shifted to increasing extensions (**Figure 6.a**). At large pulling rates we observed a similar response, although the peak decrement and peak position shift were more gradual (**Figure 6.b**). In the presence of 1 M NaCl the rightward shift of the peak was completely alleviated (**Figure 6.c**), but increasing TMPyP concentrations continued to reduce the peak value.

**Figure 5.**
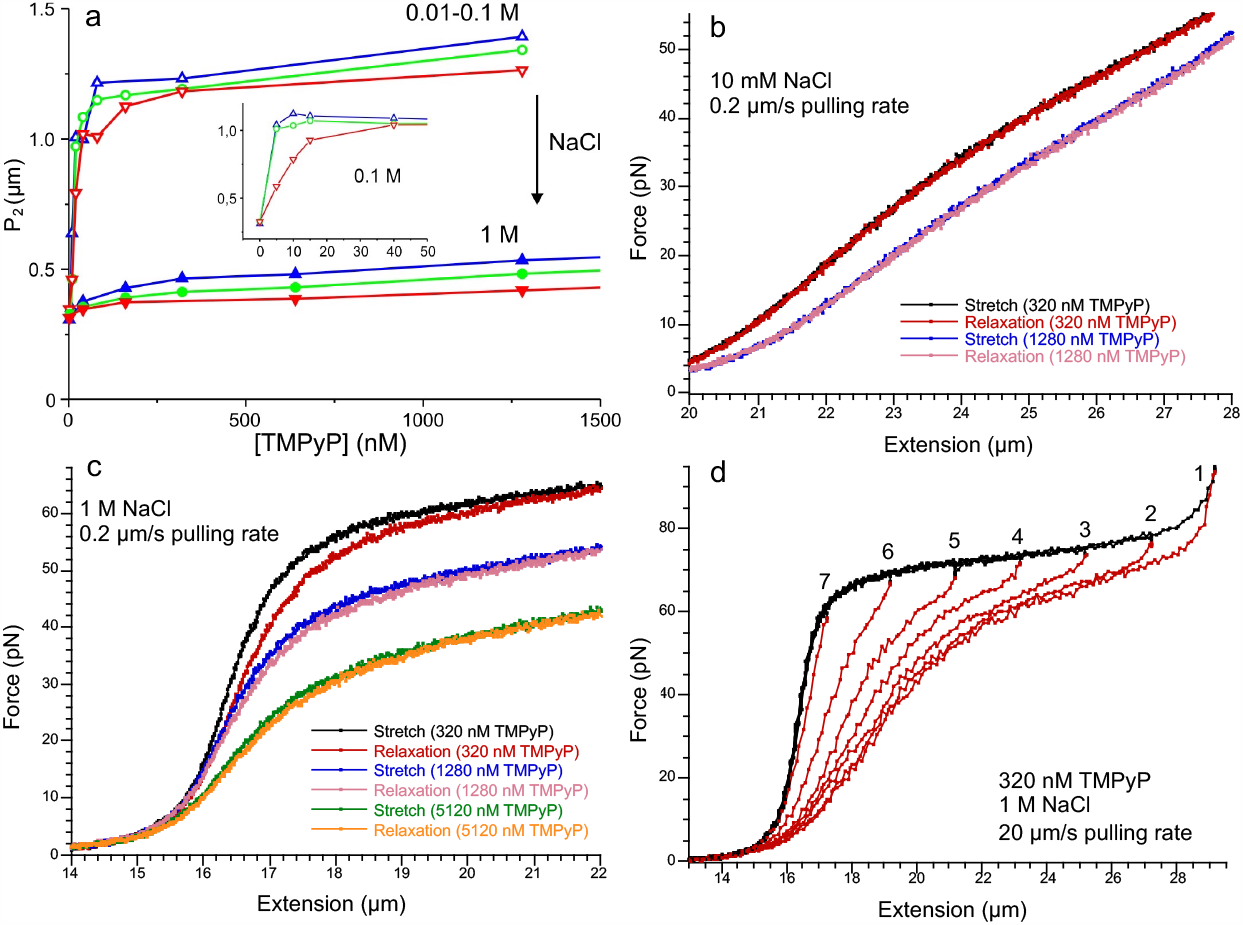
Effect of TMPyP on dsDNA nanomechanics in the enthalpic regime. **a)** P_2_ (sigmoid step width) as a function of TMPyP concentration, pulling rate and NaCl concentration (0.01, 0.1 and 1M). The pulling rates of 0.2 μm/s, 2 μm/s, 20 μm/s are indicated with blue, green and red, respectively. 0.01-0.1 M indicates that the parameters in these two cases are close to each other within error. **Inset**, low-TMPyP-concentration (<50 nM) regime of the 0.1 M NaCl data. Analysis of mechanical reversibility at 0.2 μm/s pulling rate in the presence of 10 mM NaCl and TMPyP concentrations indicated in the legend. The plot focuses on the enthalpic region. **c)** Analysis of mechanical reversibility at 0.2 μm/s pulling rate in the presence of 1 M NaCl and TMPyP concentrations indicated in the legend. **d)** Effect of maximal stretch on mechanical reversibility at 20 μm/s pulling rate in the presence of 1 M NaCl and 320 nM TMPyP. The DNA molecule was stretched and relaxed in consecutive mechanical cycles with progressively decreasing maximal stretch length. The maximum stretch lengths are indicated with numbers above the force curves. The video of this experiment (**Supplementary Video**) is shown in the **Supplementary Information**.

**Figure 6.**
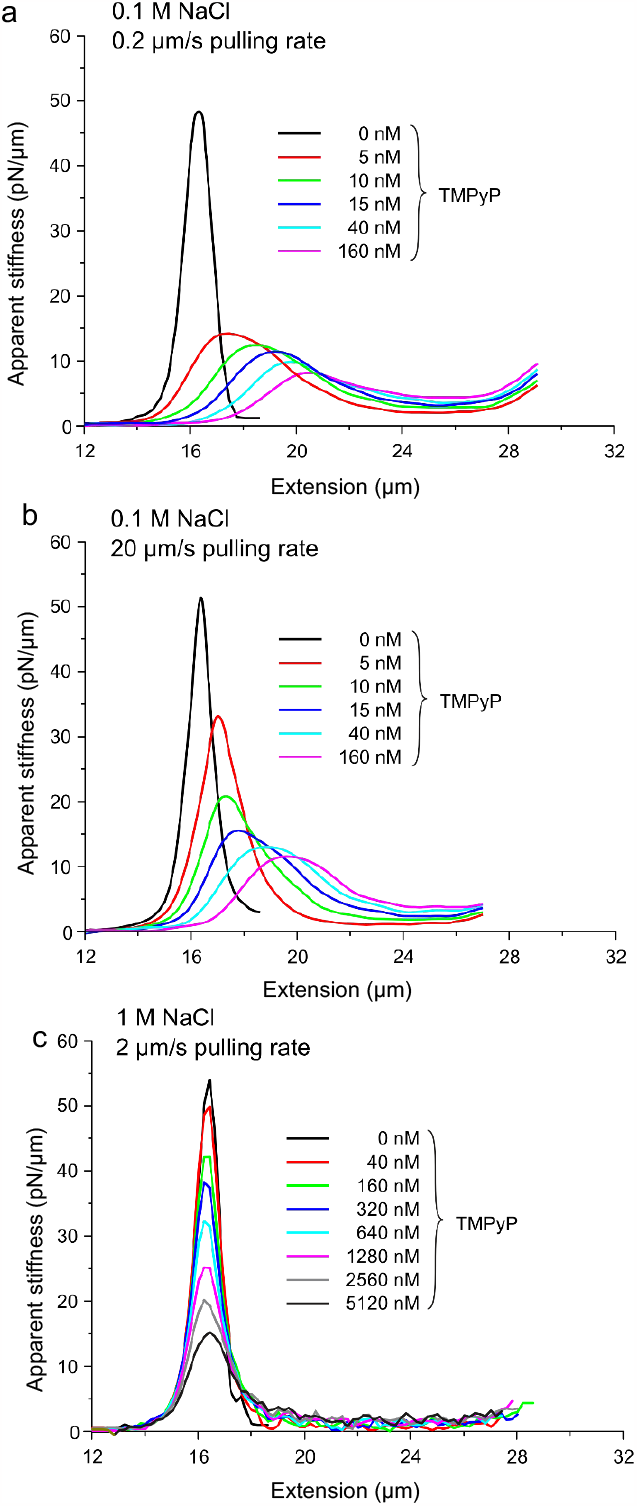
Instantaneous apparent stiffness of DNA generated by numerical derivation of stretch force-extension curves. Three series of TMPyP-dependent curves are shown: **a)** at 0.1 M NaCl and 0.2 μm/s pulling speed, **b)** at 0.1 M NaCl and 20 μm/s and **c)** at 1 M NaCl and 2 μm/s. In the first two series (a and b) smoothing was applied, and in the third (c) the raw, unsmoothed data are shown. The colors correspond to the different TMPyP concentrations indicated in the legend.

### Effect of TMPyP on the overstretch transition

The overstretch transition is characterized by its slope and height, which are reflected in the *P*_3_ and *P*_4_ parameters of the fitting function, respectively. *P*_3_ increased rapidly, then slowly, as a function of TMPyP concentration (**Figure 7.a**). The effect was significantly reduced, but it was not completely alleviated, upon increasing NaCl concentration to 1 M. At 1 M NaCl the pulling rate-dependence increased, suggesting that the molecular system was shifted away from equilibrium. To test for this possibility, we compared the stretch and relaxation force curves at high (20 μm/s) pulling rates (**Figure 7.b**). Indeed, a force hysteresis was present (black and red curves in **Figure 7.b**), which progressively increased with increasing the extension across the overstretch transition. By contrast, at low NaCl (10 mM) the hysteresis was minimal (blue and pink curves).

**Figure 7.**
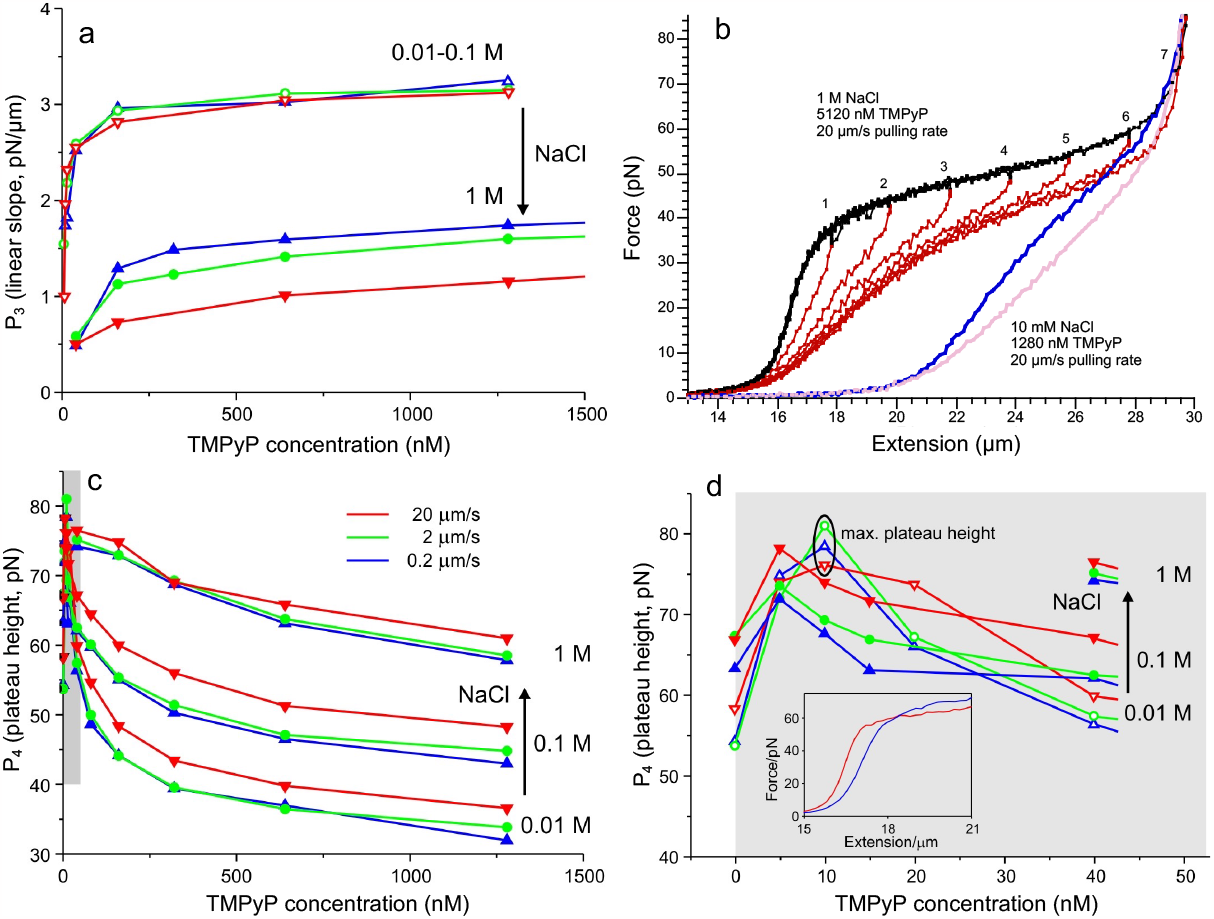
Effect of TMPyP on DNA nanomechanics in the overstretch plateau regime. **a)** P_3_ (linear slope) as a function of TMPyP concentration, pulling rate and NaCl concentration (0.01, 0.1 and 1M). The pulling rates of 0.2 μm/s, 2 μm/s, 20 μm/s are indicated with blue, green and red, respectively. 0.01-0.1 M indicates that the parameters in these two cases are close to each other within error. **b)** Analysis of mechanical reversibility at 20 μm/s pulling rate in two different experimental conditions: 1 M NaCl and 5120 nM TMPyP, stretch and relaxation indicated in black and red, respectively; 10 mM NaCl and 1280 nM TMPyP, stretch and relaxation indicated in blue and pink, respectively. In the 1 M NaCl experiment the DNA molecule was stretched and relaxed in consecutive mechanical cycles with progressively increasing maximal stretch length. The maximum stretch lengths are indicated with numbers above the force curves. **c)** P_4_ (plateau height), measured at an extension of 25 μm, as a function of TMPyP concentration, pulling rate and NaCl concentration (0.01, 0.1 and 1M). **d)** Expanded view of **c)** in the TMPyP concentration range of 0-50 nM. Plateau height peaks at a TMPyP concentration of 10 nM. **Inset**, force-extension curves at low TMPyP concentrations (10 mM NaCl, 0.2 μm/s pulling rate) shown to demonstrate the local peaking effect of TMPyP on the plateau height. Red and blue correspond to 5 and 10 nM TMPyP, respectively.

*P*_4_, the height of the overstretch transition, measured systematically at the arbitrarily assigned extension of 25 μm, decreased progressively as function of TMPyP concentration (**Figure 7.c**). Increasing NaCl concentration gradually reduced this effect. We also observed pulling rate-dependence, which concurs with its effect on the *P*_3_ parameter (see **Figure 7.b**). Even though *P*_4_ decreased with increasing TMPyP concentration, below a TMPyP concentration of 10 nM we observed a transient increment and local maximum (**Figure 7.d**). The observation is substantiated by the crossing of the FECs in the overstretch transition (**Figure 7.d inset**).

### Effect of TMPyP on the asymptotic regime

*P*_5_, which reflects the maximal length of the overstretched DNA molecule, increased gradually towards a maximum as a function of TMPyP concentration (**Figure 8.a**). In 1 M NaCl, *P*_5_ was reduced considerably at low TMPyP concentration, then it increased apparently towards the same maximum value. Pulling rate had minimal effect on this parameter and on its NaCl concentration dependence.

**Figure 8.**
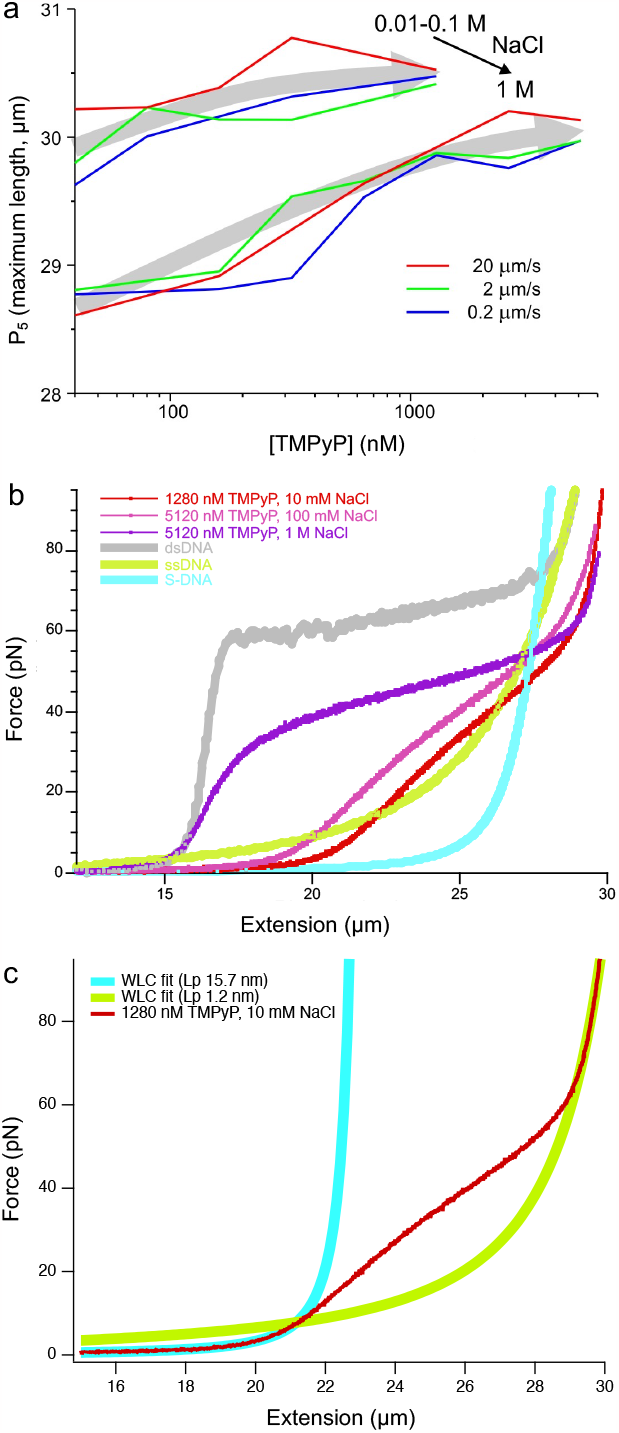
Effect of TMPyP on DNA nanomechanics in the asymptotic regime. **a)** P_5_ (maximum molecular length) as a function of TMPyP concentration, pulling rate and NaCl concentration. 0.01-0.1 M means that the parameters in these two cases are close to each other within error. Gray arrows indicate the trend that P_5_ reaches a plateau. **b)** Comparison of canonical, theoretical and extreme (maximum tested TMPyP concentrations) experimental force-extension curves of DNA. The experimental FECs are equilibrium traces obtained at low (0.2 μm/s) pulling rates. Equilibrium FEC (red trace) of DNA obtained at 1280 nM TMPyP, 10 mM NaCl and 0.2 μm/s pulling rate with wormlike-chain (WLC) fits on the entropic (<7 pN, light blue) and asymptotic (>70 pN, light green) regimes.

## Discussion

### TMPyP binds to dsDNA by two modes and elongates it

In the present work we investigated the effect of TMPyP, a chemical widely used in photodynamic therapy (25,27,32,52-57) and G-quadruple stabilization (40,58-64), on DNA nanomechanics. Even though the interaction of TMPyP with DNA has been investigated extensively, how it might alter the nanomechanical behavior of DNA remained unknown. Because many important DNA-binding proteins are mechanoenzymes (e.g., DNA-and RNA-polymerases, etc.), understanding DNA’s nanomechanical response to pharmacological perturbations is of great importance.

The addition of TMPyP to dsDNA in increasing concentration resulted in a complex array of changes (**Figures 2, S1**) with respect to DNA’s canonical force *versus* extension curve (FEC) (**Figure 1**). Moreover, adding NaCl to the molecular system, which is commonly used for stabilizing the double-helical structure and for electrostatic screening (4,6), resulted in further, complex changes in the FEC. Because in nanomechanical experiments force is used to distort molecular structure as a function of time, and as a result the equilibrium of the binding reaction may be constantly shifted, the thermodynamic state of the system is an important question. On one hand, it may be desired to characterize DNA in a chemically constant system, in which the ratio of the reactant to DNA in the complex remains steady (6). We found that the binding of TMPyP to DNA is very fast on realistic time scales of nanomechanical experiments (data not shown), which precluded the characterization of a molecular system in which the ratio of DNA-bound TMPyP molecules remained constant. On the other hand, it may be desired to characterize DNA in thermodynamic equilibrium, in which the TMPyP-DNA complex is in chemical and conformation equilibrium throughout the mechanical stretch-relaxation cycle. Such an equilibrium is characterized by the absence of force hysteresis (65). Increasing the pulling rate pushes the molecular system away from equilibrium, and *vice versa*. To test for equilibrium, we therefore exposed the TMPyP-DNA complex to different pulling rates, which resulted in a large, multi-dimensional dataset (**Figure S1**).

To sort between the types of effects TMPyP and NaCl may have on DNA, we introduced an empirical mathematical model with which we fitted the FECs (**Figure 3**). The significance of this simple model lies in the fact that its parameters reflect the physical characteristics of DNA. Accordingly, we were able to follow changes in the contour length and compliance of dsDNA, the average force and cooperativity of the overstretch transition, and the maximal length of overstretched DNA.

TMPyP caused a significant, up to 37% increase in the contour length of dsDNA, reflected in the change of the *P*_1_ parameter (**Figure 4.a**), in an essentially pulling rate-independent manner, indicating that the structural changes caused by TMPyP take place rapidly. Most of the lengthening occurs in the low TMPyP concentration range (<40 nM), followed by a more gradual TMPyP concentration-dependent extension, suggesting that TMPyP binds in at least two binding modes (42,44). The length increment could be completely inhibited by 1 M NaCl, indicating that the mechanism of TMPyP binding to DNA is electrostatic. It has been shown before that TMPyP intercalates into DNA (66-69), and that intercalators cause DNA lengthening (6,8,10,12,13,48-51). Thus, we conclude that the primary length increment is caused by TMPyP intercalation. It has been also found that TMPyP binds to DNA with alternative mechanisms: major and minor groove binding (67,70,71). Most plausibly, these binding mechanisms altogether lead to the secondary DNA lengthening step. To dissect the binding mechanisms further, we measured the length increment at given forces in an analysis employed before in the investigation of intercalator-DNA interactions (6). The force-dependent length change could be fitted with mono-exponential functions (**Figure 4.b**) that ran essentially parallel with each other, indicating that the force-dependent TMPyP binding kinetics are identical, regardless of TMPyP concentration. Thus, force simply makes room in dsDNA for further TMPyP binding, but otherwise the binding reaction is unaltered. Extrapolation to zero force allowed us to calculate the length change of dsDNA induced by TMPyP binding in mechanically relaxed conditions, which supports prior observations obtained in ensemble measurements (36,37,56). The TMPyP concentration-dependent double-exponential functions of length change (**Figure 4.c**) also appeared similar by large. However, the fast and slow apparent rate constants of the double-exponential function decreased and increased as a function of force (**Figure S2.b**) in the investigated range, suggesting that the mechanical state of DNA may actually impose slight changes in the binding reaction. Unveiling the mechanisms behind this observation await further experimental investigation.

### TMPyP binding increases apparent dsDNA compliance

TMPyP caused a large, nearly step-like increase in dsDNA compliance within a narrow concentration range (0-10 nM), as judged from the change in the *P*_2_ parameter (**Figure 5.a**). The slight pulling rate-dependence of the *P*_2_ parameter suggested that the the molecular system may not be in equilibrium. The DNA-softening effect of TMPyP could be almost completely reversed by raising the NaCl concentration to 1 M. Because upon stretching DNA with force further and further, TMPyP molecules keep binding (see **Figure 4.b**), the system is in progressive chemical change. Therefore, the molecular system is not a true elastic body, and viscous behavior may arise depending on the kinetic and thermodynamic state of the TMPyP-and Na-DNA binding reactions. We tested for thermodynamic equilibrium, and found that it prevails as long as DNA is prevented from entering the overstretch transition (**Figure 5.d**). Interestingly, NaCl has a differential effect on dsDNA in this region of the FEC: whereas the original contour length of dsDNA is completely recovered, the softening effect of TMPyP persists even at high (1 M) NaCl concentration, suggesting that NaCl competes differently with TMPyP binding, depending on the binding mechanism. Altogether, dsDNA behaves as an apparent elastic body, the stiffness of which can be sensitively modulated by TMPyP (in the nM concentration range) and by ionic strength.

### TMPyP binding reduces the cooperativity and overall force of overstretch transition

Upon reaching a threshold force, typically ∼60 pN, under conditions resembling physiological, dsDNA goes through a cooperative overstretch transition characterized by a significant (>60%) lengthening that occurs within a narrow (∼15 pN) force range (4). Three main processes are thought to occur during this transition, the ratio of which are influenced by the number of nicks along DNA and environmental parameters such as ionic strength (72,73): conversion of B-DNA to S-DNA, melting bubble formation and strand unpeeling. Because these processes are affected by the strength of association between the DNA strands, the average height is thought to reflect the stability of the double-stranded DNA structure. The cooperativity of the transition is manifested in the narrow force range, hence the low slope of the FEC in this regime, and is related to the processes running linearly along the contour of the DNA chain. We found that the overstretch transition was greatly altered by addition of TMPyP and then NaCl (**Figure 7**). Cooperativity, reflected in the inverse of the *P*_3_ parameter, was significantly reduced within a narrow TMPyP range (0-40 nM), then continued to decrease as a function of increasing TMPyP concentration (**Figure 7.a**). Conceivably, the TMPyP molecules that bound to dsDNA remain attached throughout the transition and act as road-blocks that inhibit the progression of the molecular changes along the chain. Increasing NaCl concentration to 1 M partially restored cooperativity, and a strong pulling-rate dependence was present. Considering that 1 M NaCl restores the contour length of dsDNA completely (**Figure 4.a**), in the presence of high TMPyP and NaCl concentrations the roadblock TMPyP molecules are likely ones that bind newly to DNA during the overstretch transition, plausibly to ssDNA regions. NaCl competes inefficiently with these newly bound roadblock TMPyP molecules which also inhibit the re-formation of dsDNA, resulting in a marked force hysteresis (**Figure 7.b**, black and red traces). At high TMPyP but low NaCl concentrations (**Figure 7.b**, blue and pink traces) a lenghtened dsDNA and very little hysteresis are observed, which raises the possibility that the intercalated/groove-bound TMPyP population can be converted directly into the ssDNA-bound one.

The average force of the overstretch transition, reflected in the *P*_4_ parameter, decreased progressively as a function of increasing TMPyP concentration, and the effect was partially restored by increasing the concentration of NaCl (**Figure 7.c**). Interestingly, however, the *P*4 decrease was not monotonic, but a local maximum was observed at 10 nM TMPyP (**Figure 7.d**). This surprising finding suggests that the local chemical equilibria of the different binding modes become re-arranged in between the reactions stabilizing and de-stabilizing dsDNA. Conceivably, in the low TMPyP concentration range the stabilizing effects of intercalation and groove binding dominate (36,37,42), whereas at higher TMPyP concentrations the de-stabilizing effects of ssDNA binding become overwhelming. In support, the interaction with ssDNA-binding proteins results in similar FECs (13,22,23). Furthermore, it has been shown that the binding of actinomycin D (ActD) to DNA, which may occur *via* intercalation between dsDNA base pairs (74-78), ssDNA association (79-82) and ssDNA base intercalation (83,84), results in the lowering of the average overstretch force and reduction of cooperativity (85). Altogether, the complex array of TMPyP-and NaCl-induced effects on the overstretch transition of DNA is determined by a shift between the stabilizing and de-stabilizing TMPyP-DNA binding modes, and by the differential screening of intercalating and non-intercalating TMPyP-DNA association by NaCl.

### TMPyP binds to ssDNA and elongates it

Upon reaching extreme stretch, dsDNA is eventually converted into ssDNA, in which the strands are held together by a small number of hydrogen bonds. Thus, in the asymptotic regime (**Figure 1**) the FEC is set by the properties (contour and persistence lengths) of ssDNA. Our observations on the effects of TMPyP on the overstretch stransition already raised the possibility that TMPyP is able to bind ssDNA directly (**Figure 7**). Upon the addition of TMPyP in increasing concentrations, the contour length of ssDNA, reflected in the *P*_5_ parameter, increased gradually towards a maximum (**Figure 8.a**). At a NaCl concentration of 1 M, the contour length increment towards the same maximum was more pronounced. In other words, at high concentrations of NaCl ssDNA is contracted, and TMPyP competes with NaCl, leading to the lengthening of the molecule. The NaCl-induced contraction is likely caused by a decrease in the electrostatic persistence length of ssDNA due to electrostatic screening by Na+ ions. The competing effect by TMPyP is then plausibly caused by the intercalation of the positively charged molecules in between the bases of ssDNA. Altogether, the binding of TMPyP at high concentrations results in the lengthening of ssDNA by >3%.

### Possible structure of the TMPyP-bound DNA molecule

TMPyP binding in different modes combined with mechanical force converts DNA into a yet unknown structure. To estimate the structure, we compared the FEC of DNA in high TMPyP concentration with the extreme scenarios of the control dsDNA, ssDNA and S-DNA (**Figure 8.b**). At low (10-100 mM) NaCl and high (>1280 nM) TMPyP concentrations a highly extended and compliant dsDNA is seen, which is converted by a non-cooperative force-driven transition into a structure that is longer then than the control ssDNA. 1 M NaCl restores the contour length and some of the stiffness of dsDNA, which is converted by a more-or-less cooperative transition into a similarly overstretched structure. We exclude the possibility that any part of the length change might be due to G-quadruplexes, which are known to be stabilized by TMPyP (39,40,62,86). Even though λ-phage DNA is predicted to contain 30 G-quadruplex sequences (**Figure S4**), their formation would require *a priori* dsDNA denaturation. Furthermore, the mechanically-driven rupture of G-quadruplexes is expected to result in sawteeth at low forces (87,88), which we have not observed in the FECs. To investigate the nature of the TMPyP-bound DNA further, we fitted the low-(<7 pN) and high-force (>70 pN) sections of the FEC with the inextensible wormlike-chain (WLC) model (equation 5) (46) (**Figure 8.c**). According to the calculated persistence lengths (*L*_P_), at low and high forces the TMPyP-bound DNA resembles an S-DNA (*L*_P_ = 15.7 nm) and an ssDNA (*L*_P_ = 1.2 nm), respectively. Thus, the TMPyP-loaded dsDNA carries similarities to S-DNA, as it has a reduced bending rigidity, and it is extended due to the wedging of TMPyP molecules in between the base pairs and a probable partial unwinding of the double helix.

Finally, it is worth pointing out that the employed TMPyP concentrations fall below or well within those used in *in vitro* photodynamic therapy (52,55,89). Therefore, the structural and nanomechanical changes documented here are highly relevant during the therapeutic applications of TMPyP. Furthermore, the largest amplitude of the nanomechanical changes in DNA are evoked in a relatively narrow nanomolar (5-40 nM) TMPyP concentration range. Thus, an interplay between nanomolar TMPyP concentrations and piconewton forces may tune DNA’s structural and nanomechanical characteristics, thereby controlling the myriad of DNA-associated mechanoenzymatic processes.

## Conclusions

Here we have uncovered a complex array of TMPyP-induced nanomechanical changes in DNA. TMPyP binds to dsDNA with two mechanisms: intercalation and groove binding. Intercalation is a fast process leading to dsDNA lengthening, whereas groove binding is an order of magnitude slower. Force enhances TMPyP binding, but the kinetic properties of the reaction are only slightly altered. DNA extension simply makes room for the further binding of TMPyP molecules. TMPyP binding reduces the apparent instantaneous stiffness of dsDNA. TMPyP initially (at low concentrations) stabilizes, then (at high concentrations) destabilizes dsDNA. The cooperativity of the overstretch transition is reduced due to road-blocks that slow the transition. TMPyP binds, or stays bound to ssDNA, thereby lengthening it. ssDNA-bound TMPyP inhibits the re-formation of dsDNA. NaCl efficiently competes with TMPyP for DNA binding, but differentiates between the intercalation and groove-binding processes. As a result, the dsDNA contour length is efficiently recovered, but the rest of the TMPyP effects partially remain. At high NaCl concentrations, ssDNA contour length is reduced, most likely due to electrostatic screening, and TMPyP competes with NaCl by screening its length-reducing effect. The complex, TMPyP concentration-dependent changes in DNA nanomechanics provide a wide array of possibilities to modulate the force-dependent processes within the genome and may have significant therapeutic implications.

## Abbreviations

TMPyP: tetrakis(4-N-methyl)pyridyl-porphyrin;
FEC: force versus extension curve;
ds: double stranded;
ss: single stranded;
WLC: wormlike chain

## Data Availability

All experimental data are available upon request by writing to the corresponding author.

## Author Contributions

Conceptualization, M.K. and B.K.; methodology, B.K.; software, L.H. and B.K.; validation, B.K., M.K. and L.H.; formal analysis, L.H. and B. K.; investigation, B.K. and E.S.; resources, M.K.; data curation, X.X.; writing—original draft preparation, B.K.; writing—review and editing, M.K., L. H., G. Cs., H. T., B. K., A. O. and B. K.; visualization,

L.H. and B.K.; supervision, M.K.; project administration, M.K.; funding acquisition, M.K. All authors have read and agreed to the published version of the manuscript.

## Funding

This research was funded by the ÚNKP-21-3-II-SE-37 New National Excellence Program of The Ministry for Innovation and Technology and by Predoctoral Scholarship awarded by the Doctoral School of Semmelweis University to B.K. Grants from the Hungarian National Research, Development and Innovation Office (K135360 to M.K., Project no. NVKP_16-1–2016-0017 ‘National Heart Program’, and the 2020-1.1.6-JÖVŐ-2021-00013 grant), the Ministry for Innovation and Technology of Hungary (Thematic Excellence Programme 2020-4.1.1.-TKP2020 within the framework of the Therapeutic Development and Bioimaging thematic programs of Semmelweis University; TKP2021-NVA-15 and TKP2021-EGA-23 which have been implemented from the National Research, Development and Innovation Fund, financed under the TKP2021-NVA and TKP2021-EGA funding schemes, respectively), and the European Union (Project no. RRF-2.3.1-21-2022-00003).

## Institutional Review Board Statement

Not applicable.

## Informed Consent Statement

Not applicable.

## Data Availability Statement

Not applicable.

## Acknowledgments

We thank Dávid Szöllősi for his help with data animation, Mónika Komárné Drabbant, Krisztina Lór and Zsófia Kovács for technical assistance, and Erzsébet Suhajda for reviewing the manuscript and providing insightful thoughts.

## Conflicts of Interest

The authors declare no conflict of interest.

## Notes

### Competing Interest Statement

The authors have declared no competing interest.

## References

1. Gross, P., Laurens, N., Oddershede, L.B., Bockelmann, U., Peterman, E.J.G. and Wuite, G.J.L. (2011) Quantifying how DNA stretches, melts and changes twist under tension. Nature Physics, 7, 731–736.

2. Harlepp, S., Chardon, E., Bouché, M., Dahm, G., Maaloum, M. and Bellemin-Laponnaz, S. (2019) N-Heterocyclic Carbene-Platinum Complexes Featuring an Anthracenyl Moiety: Anti-Cancer Activity and DNA Interaction. IJMS, 20, 4198.

3. Kaczorowska, A., Lamperska, W., Frączkowska, K., Masajada, J., Drobczyński, S., Sobas, M., Wróbel, T., Chybicka, K., Tarkowski, R., Kraszewski, S. et al. (2020) Profound Nanoscale Structural and Biomechanical Changes in DNA Helix upon Treatment with Anthracycline Drugs. IJMS, 21, 4142.

4. Smith, S.B., Cui, Y. and Bustamante, C. (1996) Overstretching B-DNA: The Elastic Response of Individual Double-Stranded and Single-Stranded DNA Molecules. Science, 271, 795–799.

5. van Mameren, J., Gross, P., Farge, G., Hooijman, P., Modesti, M., Falkenberg, M., Wuite, G.J.L. and Peterman, E.J.G. (2009) Unraveling the structure of DNA during overstretching by using multicolor, single-molecule fluorescence imaging. Proc. Natl. Acad. Sci. U.S.A., 106, 18231–18236.

6. Biebricher, A.S., Heller, I., Roijmans, R.F., Hoekstra, T.P., Peterman, E.J. and Wuite, G.J. (2015) The impact of DNA intercalators on DNA and DNA-processing enzymes elucidated through force-dependent binding kinetics. Nat Commun, 6, 7304.

7. Renger, R., Morin, J.A., Lemaitre, R., Ruer-Gruss, M., Jülicher, F., Hermann, A. and Grill, S.W. (2022) Co-condensation of proteins with single- and double-stranded DNA. Proc. Natl. Acad. Sci. U.S.A., 119, e2107871119.

8. Kretzer, B., Kiss, B., Tordai, H., Csík, G., Herényi, L. and Kellermayer, M. (2020) Single-Molecule Mechanics in Ligand Concentration Gradient. Micromachines, 11, 212.

9. (2012) Methods for studying nucleic acid/drug interactions. CRC Press, Boca Raton.

10. Almaqwashi, A.A., Paramanathan, T., Rouzina, I. and Williams, M.C. (2016) Mechanisms of small molecule–DNA interactions probed by single-molecule force spectroscopy. Nucleic Acids Research, 44, 3971–3988.

11. Halma, M.T.J., Tuszynski, J.A. and Wuite, G.J.L. (2022) Optical tweezers for drug discovery. Drug Discovery Today, 103443.

12. McCauley, M.J. and Williams, M.C. (2007) Mechanisms of DNA binding determined in optical tweezers experiments. Biopolymers, 85, 154–168.

13. McCauley, M.J. and Williams, M.C. (2009) Optical tweezers experiments resolve distinct modes of DNA-protein binding. Biopolymers, 91, 265–282.

14. Amanzadeh, E., Mohabatkar, H. and Biria, D. (2014) Classification of DNA Minor and Major Grooves Binding Proteins According to the NLSs by Data Analysis Methods. Appl Biochem Biotechnol, 174, 437–451.

15. Chaurasiya, K.R., Paramanathan, T., McCauley, M.J. and Williams, M.C. (2010) Biophysical characterization of DNA binding from single molecule force measurements. Physics of Life Reviews, 7, 299–341.

16. Husale, S., Grange, W. and Hegner, M. (2002) DNA Mechanics Affected by Small DNA Interacting Ligands. Single Mol., 3, 91–96.

17. Krautbauer, R., Fischerländer, S., Allen, S. and Gaub, H.E. (2002) Mechanical Fingerprints of DNA Drug Complexes. Single Mol., 3, 97–103.

18. Krautbauer, R., Pope, L.H., Schrader, T.E., Allen, S. and Gaub, H.E. (2002) Discriminating small molecule DNA binding modes by single molecule force spectroscopy. FEBS Letters, 510, 154–158.

19. Mihailovic, A., Vladescu, I., McCauley, M., Ly, E., Williams, M.C., Spain, E.M. and Nuñez, M.E. (2006) Exploring the Interaction of Ruthenium(II) Polypyridyl Complexes with DNA Using Single-Molecule Techniques. Langmuir, 22, 4699–4709.

20. Sischka, A., Toensing, K., Eckel, R., Wilking, S.D., Sewald, N., Ros, R. and Anselmetti, D. (2005) Molecular Mechanisms and Kinetics between DNA and DNA Binding Ligands. Biophysical Journal, 88, 404–411.

21. Stassi, S., Marini, M., Allione, M., Lopatin, S., Marson, D., Laurini, E., Pricl, S., Pirri, C.F., Ricciardi, C. and Di Fabrizio, E. (2019) Nanomechanical DNA resonators for sensing and structural analysis of DNA-ligand complexes. Nature Communications, 10, 1690.

22. Shokri, L., Marintcheva, B., Richardson, C.C., Rouzina, I. and Williams, M.C. (2006) Single Molecule Force Spectroscopy of Salt-dependent Bacteriophage T7 Gene 2.5 Protein Binding to Single-stranded DNA. Journal of Biological Chemistry, 281, 38689–38696.

23. Shokri, L., Rouzina, I. and Williams, M.C. (2009) Interaction of bacteriophage T4 and T7 single-stranded DNA-binding proteins with DNA. Phys. Biol., 6, 025002.

24. Garcia-Sampedro, A., Tabero, A., Mahamed, I. and Acedo, P. (2019) Multimodal use of the porphyrin TMPyP: From cancer therapy to antimicrobial applications. J. Porphyrins Phthalocyanines, 23, 11–27.

25. Lang, K., Mosinger, J. and Wagnerová, D.M. (2004) Photophysical properties of porphyrinoid sensitizers non-covalently bound to host molecules; models for photodynamic therapy. Coordination Chemistry Reviews, 248, 321–350.

26. Tada-Oikawa, S., Oikawa, S., Hirayama, J., Hirakawa, K. and Kawanishi, S. (2009) DNA Damage and Apoptosis Induced by Photosensitization of 5,10,15,20-Tetrakis (<i>N</i> -methyl-4-pyridyl)-21 <i>H</i>, 23 <i>H</i> -porphyrin <i>via</i> Singlet Oxygen Generation. Photochemistry and Photobiology, 85, 1391–1399.

27. Villanueva, A., Stockert, J.C., Cañete, M. and Acedo, P. (2010) A new protocol in photodynamic therapy: enhanced tumour cell death by combining two different photosensitizers. Photochem Photobiol Sci, 9, 295–297.

28. Kakiuchi, T., Ito, F. and Nagamura, T. (2008) Time-resolved studies of energy transfer from meso-tetrakis(N-methylpyridinium-4-yl)-porphyrin to 3,3’-diethyl-2,2’-thiatricarbocyanine iodide along deoxyribonucleic acid Chain. J Phys Chem B, 112, 3931–3937.

29. Mutsamwira, S., Ainscough, E.W., Partridge, A.C., Derrick, P.J. and Filichev, V.V. (2014) DNA duplex as a scaffold for a ground state complex formation between a zinc cationic porphyrin and phenylethynylpyren-1-yl. Journal of Photochemistry and Photobiology A: Chemistry, 288, 76–81.

30. Pathak, P., Yao, W., Hook, K.D., Vik, R., Winnerdy, F.R., Brown, J.Q., Gibb, B.C., Pursell, Z.F., Phan, A.T. and Jayawickramarajah, J. (2019) Bright G-Quadruplex Nanostructures Functionalized with Porphyrin Lanterns. J Am Chem Soc, 141, 12582–12591.

31. Stulz, E. (2017) Nanoarchitectonics with Porphyrin Functionalized DNA. Acc. Chem. Res., 50, 823–831.

32. Diogo, P., Fernandes, C., Caramelo, F., Mota, M., Miranda, I.M., Faustino, M.A.F., Neves, M.G.P.M.S., Uliana, M.P., de Oliveira, K.T., Santos, J.M. et al. (2017) Antimicrobial Photodynamic Therapy against Endodontic Enterococcus faecalis and Candida albicans Mono and Mixed Biofilms in the Presence of Photosensitizers: A Comparative Study with Classical Endodontic Irrigants. Front. Microbiol., 8.

33. Gonzales, F.P., Felgenträger, A., Bäumler, W. and Maisch, T. (2013) Fungicidal photodynamic effect of a twofold positively charged porphyrin against <i>Candida albicans</i> planktonic cells and biofilms. Future Microbiology, 8, 785–797.

34. Grinholc, M., Rodziewicz, A., Forys, K., Rapacka-Zdonczyk, A., Kawiak, A., Domachowska, A., Golunski, G., Wolz, C., Mesak, L., Becker, K. et al. (2015) Fine-tuning recA expression in Staphylococcus aureus for antimicrobial photoinactivation: importance of photo-induced DNA damage in the photoinactivation mechanism. Appl Microbiol Biotechnol, 99, 9161–9176.

35. Quiroga, E.D., Alvarez, M.G. and Durantini, E.N. (2010) Susceptibility of Candida albicans to photodynamic action of 5,10,15,20-tetra(4-N-methylpyridyl)porphyrin in different media: Photodynamic action of TMPyP on C. albicans. FEMS Immunology & Medical Microbiology, 60, 123–131.

36. Zupán, K., Herényi, L., Tóth, K., Egyeki, M. and Csík, G. (2005) Binding of Cationic Porphyrin to Isolated DNA and Nucleoprotein Complex: Quantitative Analysis of Binding Forms under Various Experimental Conditions. Biochemistry, 44, 15000–15006.

37. Zupán, K., Herényi, L., Tóth, K., Majer, Z. and Csík, G. (2004) Binding of Cationic Porphyrin to Isolated and Encapsidated Viral DNA Analyzed by Comprehensive Spectroscopic Methods. Biochemistry, 43, 9151–9159.

38. Kim, M.-Y., Gleason-Guzman, M., Izbicka, E., Nishioka, D. and Hurley, L.H. (2003) The Different Biological Effects of Telomestatin and TMPyP4 Can Be Attributed to Their Selectivity for Interaction with Intramolecular or Intermolecular G-Quadruplex Structures. Cancer Res., 63, 3247–3256.

39. Molnar, O.R., Vegh, A., Somkuti, J. and Smeller, L. (2021) Characterization of a G-quadruplex from hepatitis B virus and its stabilization by binding TMPyP4, BRACO19 and PhenDC3. Sci Rep, 11, 23243.

40. Ruan, T.L., Davis, S.J., Powell, B.M., Harbeck, C.P., Habdas, J., Habdas, P. and Yatsunyk, L.A. (2017) Lowering the overall charge on TMPyP4 improves its selectivity for G-quadruplex DNA. Biochimie, 132, 121–130.

41. Taka, T., Joonlasak, K., Huang, L., Randall Lee, T., Chang, S.-W.T. and Tuntiwechapikul, W. (2012) Down-regulation of the human VEGF gene expression by perylene monoimide derivatives. Bioorganic & Medicinal Chemistry Letters, 22, 518–522.

42. Fiel, R.J. (1989) Porphyrin—Nucleic Acid Interactions: A Review. Journal of Biomolecular Structure and Dynamics, 6, 1259–1274.

43. Bustamante, C., Gurrieri, S., Pasternack, R.F., Purrello, R. and Rizzarelli, E. (1994) Interaction of water-soluble porphyrins with single- and double-stranded polyribonucleotides. Biopolymers, 34, 1099–1104.

44. Pasternack, R.F., Brigandi, R.A., Abrams, M.J., Williams, A.P. and Gibbs, E.J. (1990) Interactions of porphyrins and metalloporphyrins with single-stranded poly(dA). Inorg. Chem., 29, 4483–4486.

45. Pasternack, R.F., Gibbs, E.J. and Villafranca, J.J. (1983) Interactions of porphyrins with nucleic acids. Biochemistry, 22, 2406–2414.

46. Bustamante, C.J., Marko, J.F., Siggia, E.D. and Smith, S.B. (1994) Entropic elasticity of lambda-phage DNA. Science, 265, 1599–1600.

47. Wenner, J.R., Williams, M.C., Rouzina, I. and Bloomfield, V.A. (2002) Salt Dependence of the Elasticity and Overstretching Transition of Single DNA Molecules. Biophysical Journal, 82, 3160–3169.

48. Almaqwashi, A.A., Zhou, W., Naufer, M.N., Riddell, I.A., Yilmaz, Ö.H., Lippard, S.J. and Williams, M.C. (2019) DNA Intercalation Facilitates Efficient DNA-Targeted Covalent Binding of Phenanthriplatin. Journal of the American Chemical Society, 141, 1537–1545.

49. Murade, C.U., Subramaniam, V., Otto, C. and Bennink, M.L. (2009) Interaction of Oxazole Yellow Dyes with DNA Studied with Hybrid Optical Tweezers and Fluorescence Microscopy. Biophysical Journal, 97, 835–843.

50. Reis, L.A., Ramos, E.B. and Rocha, M.S. (2013) DNA Interaction with Diaminobenzidine Studied with Optical Tweezers and Dynamic Light Scattering. J. Phys. Chem. B, 117, 14345–14350.

51. Vladescu, I.D., McCauley, M.J., Rouzina, I. and Williams, M.C. (2005) Mapping the Phase Diagram of Single DNA Molecule Force-Induced Melting in the Presence of Ethidium. Phys. Rev. Lett., 95, 158102.

52. Zarska, L., Mala, Z., Langova, K., Malina, L., Binder, S., Bajgar, R. and Kolarova, H. (2021) The effect of two porphyrine photosensitizers TMPyP and ZnTPPS(4) for application in photodynamic therapy of cancer cells in vitro. Photodiagnosis Photodyn Ther, 34, 102224.

53. Heffron, J., Bork, M., Mayer, B.K. and Skwor, T. (2021) A Comparison of Porphyrin Photosensitizers in Photodynamic Inactivation of RNA and DNA Bacteriophages. Viruses, 13.

54. Cenklova, V. (2017) Photodynamic therapy with TMPyP - Porphyrine induces mitotic catastrophe and microtubule disorganization in HeLa and G361 cells, a comprehensive view of the action of the photosensitizer. J Photochem Photobiol B, 173, 522–537.

55. Feese, E. and Ghiladi, R.A. (2009) Highly efficient in vitro photodynamic inactivation of Mycobacterium smegmatis. J Antimicrob Chemother, 64, 782–785.

56. Zupan, K., Egyeki, M., Toth, K., Fekete, A., Herenyi, L., Modos, K. and Csik, G. (2008) Comparison of the efficiency and the specificity of DNA-bound and free cationic porphyrin in photodynamic virus inactivation. J Photochem Photobiol B, 90, 105–112.

57. Salmon-Divon, M., Nitzan, Y. and Malik, Z. (2004) Mechanistic aspects of Escherichia coli photodynamic inactivation by cationic tetra-meso(N-methylpyridyl)porphine. Photochem Photobiol Sci, 3, 423–429.

58. Haldar, S., Zhang, Y., Xia, Y., Islam, B., Liu, S., Gervasio, F.L., Mulholland, A.J., Waller, Z.A.E., Wei, D. and Haider, S. (2022) Mechanistic Insights into the Ligand-Induced Unfolding of an RNA G-Quadruplex. J Am Chem Soc, 144, 935–950.

59. Zhou, W., Cheng, Y., Song, B., Hao, J., Miao, W., Jia, G. and Li, C. (2021) Cationic Porphyrin-Mediated G-Quadruplex DNA Oxidative Damage: Regulated by the Initial Interplay between DNA and TMPyP4. Biochemistry, 60, 3707–3713.

60. Ramos, C.I.V., Monteiro, A.R., Moura, N.M.M., Faustino, M.A.F., Trindade, T. and Neves, M. (2021) The Interactions of H(2)TMPyP, Analogues and Its Metal Complexes with DNA G-Quadruplexes-An Overview. Biomolecules, 11.

61. Molnár, O.R., Végh, A., Somkuti, J. and Smeller, L. (2021) Characterization of a G-quadruplex from hepatitis B virus and its stabilization by binding TMPyP4, BRACO19 and PhenDC3. Scientific Reports, 11, 23243.

62. Perez-Arnaiz, C., Busto, N., Santolaya, J., Leal, J.M., Barone, G. and Garcia, B. (2018) Kinetic evidence for interaction of TMPyP4 with two different G-quadruplex conformations of human telomeric DNA. Biochim Biophys Acta Gen Subj, 1862, 522–531.

63. Boschi, E., Davis, S., Taylor, S., Butterworth, A., Chirayath, L.A., Purohit, V., Siegel, L.K., Buenaventura, J., Sheriff, A.H., Jin, R. et al. (2016) Interaction of a Cationic Porphyrin and Its Metal Derivatives with G-Quadruplex DNA. J Phys Chem B, 120, 12807–12819.

64. Boncina, M., Podlipnik, C., Piantanida, I., Eilmes, J., Teulade-Fichou, M.P., Vesnaver, G. and Lah, J. (2015) Thermodynamic fingerprints of ligand binding to human telomeric G-quadruplexes. Nucleic Acids Res, 43, 10376–10386.

65. Kellermayer, M.S., Smith, S.B., Bustamante, C. and Granzier, H.L. (1998) Complete unfolding of the titin molecule under external force. J Struct Biol, 122, 197–205.

66. Le, V.H., Nagesh, N. and Lewis, E.A. (2013) Bcl-2 promoter sequence G-quadruplex interactions with three planar and non-planar cationic porphyrins: TMPyP4, TMPyP3, and TMPyP2. PLoS One, 8, e72462.

67. Lee, S., Jeon, S.H., Kim, B.J., Han, S.W., Jang, H.G. and Kim, S.K. (2001) Classification of CD and absorption spectra in the Soret band of H(2)TMPyP bound to various synthetic polynucleotides. Biophys Chem, 92, 35–45.

68. Lubitz, I., Borovok, N. and Kotlyar, A. (2007) Interaction of monomolecular G4-DNA nanowires with TMPyP: evidence for intercalation. Biochemistry, 46, 12925–12929.

69. Serra, V.V., Teixeira, R., Andrade, S.M. and Costa, S.M. (2016) Design of polyelectrolyte core-shells with DNA to control TMPyP binding. Colloids Surf B Biointerfaces, 146, 127–135.

70. Arnaud, P., Zakrzewska, K. and Meunier, B. (2003) Theoretical study of the interaction between a high-valent manganese porphyrin oxyl-(hydroxo)-Mn(IV)-TMPyP and double-stranded DNA. J Comput Chem, 24, 797–805.

71. Ishikawa, Y., Tomisugi, Y. and Uno, T. (2006) Molecular modeling of anti-parallel G-quadruplex DNA/TMPyP complexes. Nucleic Acids Symp Ser (Oxf), 331–332.

72. Bosaeus, N., El-Sagheer, A.H., Brown, T., Åkerman, B. and Nordén, B. (2014) Force-induced melting of DNA—evidence for peeling and internal melting from force spectra on short synthetic duplex sequences. Nucleic Acids Research, 42, 8083–8091.

73. King, G.A., Gross, P., Bockelmann, U., Modesti, M., Wuite, G.J.L. and Peterman, E.J.G. (2013) Revealing the competition between peeled ssDNA, melting bubbles, and S-DNA during DNA overstretching using fluorescence microscopy. Proc. Natl. Acad. Sci. U.S.A., 110, 3859–3864.

74. Lian, C., Robinson, H. and Wang, A.H.J. (1996) Structure of Actinomycin D bound with (GAAGCTTC) <sub>2</sub> and (GATGCTTC) <sub>2</sub> and Its Binding to the (CAG) <i> <sub>n</sub> </i> :(CTG) <i> <sub>n</sub> </i> Triplet Sequence As Determined by NMR Analysis. Journal of the American Chemical Society, 118, 8791–8801.

75. Müller, W. and Crothers, D.M. (1968) Studies of the binding of actinomycin and related compounds to DNA. Journal of Molecular Biology, 35, 251–290.

76. Sobell, H.M. and Jain, S.C. (1972) Stereochemistry of actinomycin binding to DNA. II. Detailed molecular model of actinomycin-DNA complex and its implications. Journal of Molecular Biology, 68, 21–34.

77. Sobell, H.M., Jain, S.C., Sakore, T.D. and Nordman, C.E. (1971) Stereochemistry of Actinomycin–DNA Binding. Nature New Biology, 231, 200–205.

78. Takusagawa, F., Dabrow, M., Neidle, S. and Berman, H.M. (1982) The structure of a pseudo intercalated complex between actinomycin and the DNA binding sequence d(GpC). Nature, 296, 466–469.

79. Chen, F.M. (2004) The nature of actinomycin D binding to d(AACCAXYG) sequence motifs. Nucleic Acids Research, 32, 271–277.

80. Rill, R.L. and Hecker, K.H. (1996) Sequence-Specific Actinomycin D Binding to Single-Stranded DNA Inhibits HIV Reverse Transcriptase and Other Polymerases. Biochemistry, 35, 3525–3533.

81. Wadkins, R.M. and Jovin, T.M. (1991) Actinomycin D and 7-aminoactinomycin D binding to single-stranded DNA. Biochemistry, 30, 9469–9478.

82. Zhou, X., Shen, Z., Li, D., He, X. and Lin, B. (2007) Study of interactions between actinomycin D and oligonucleotides by microchip electrophoresis and ESI-MS. Talanta, 72, 561–567.

83. Alexopoulos, E., Jares-Erijman, E.A., Jovin, T.M., Klement, R., Machinek, R., Sheldrick, G.M. and Usón, I. (2005) Crystal and solution structures of 7-amino-actinomycin D complexes with d(TTAGBrUT), d(TTAGTT) and d(TTTAGTTT). Acta Crystallographica Section D Biological Crystallography, 61, 407–415.

84. Wadkins, R.M., Jares-Erijman, E.A., Klement, R., Rüdiger, A. and Jovin, T.M. (1996) Actinomycin D Binding to Single-stranded DNA: Sequence Specificity and Hemi-intercalation Model from Fluorescence and1H NMR Spectroscopy. Journal of Molecular Biology, 262, 53–68.

85. Paramanathan, T., Vladescu, I., McCauley, M.J., Rouzina, I. and Williams, M.C. (2012) Force spectroscopy reveals the DNA structural dynamics that govern the slow binding of Actinomycin D. Nucleic Acids Research, 40, 4925–4932.

86. Bai, L.P., Liu, J., Han, L., Ho, H.M., Wang, R. and Jiang, Z.H. (2014) Mass spectrometric studies on effects of counter ions of TMPyP4 on binding to human telomeric DNA and RNA G-quadruplexes. Anal Bioanal Chem, 406, 5455–5463.

87. Funayama, R., Nakahara, Y., Kado, S., Tanaka, M. and Kimura, K. (2014) A single-molecule force-spectroscopic study on stabilization of G-quadruplex DNA by a telomerase inhibitor. Analyst, 139, 4037–4043.

88. Kusi-Appauh, N., Ralph, S.F., van Oijen, A.M. and Spenkelink, L.M. (2023) Understanding G-Quadruplex Biology and Stability Using Single-Molecule Techniques. J Phys Chem B, 127, 5521–5540.

89. Felgentrager, A., Gonzales, F.P., Maisch, T. and Baumler, W. (2013) Ion-induced stacking of photosensitizer molecules can remarkably affect the luminescence detection of singlet oxygen in Candida albicans cells. J Biomed Opt, 18, 045002.

