## Supplementary Information for "TMPyP binding evokes a complex, tunable nanomechanical response in DNA"

### Complex nanomechanical response of dsDNA evoked by TMPyP binding

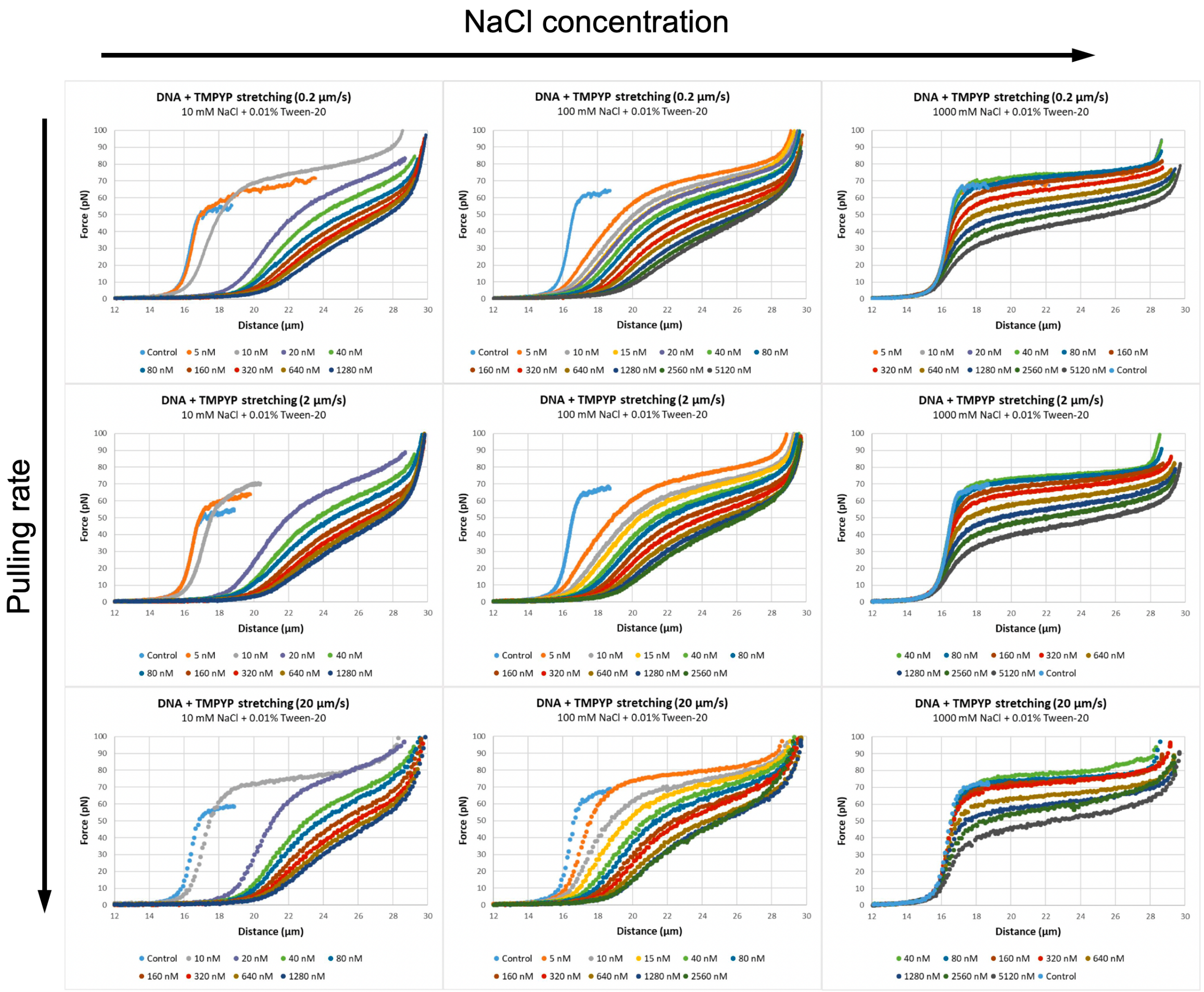


**Figure S1.** Effect of TMPyP concentration, pulling rate and NaCl concentration on the stretch force-extension curves (FEC) of λ-phage DNA. Nine sets of TMPyP concentration-dependent FECs are shown in the pulling-rate and NaCl-concentration phase spaces. The tested TMPyP concentrations are indicated below each figure. In the case of the 10 mM NaCl measurements, the maximum TMPyP concentration at which single dsDNA molecules could be reliably stretched was 1280 nM. Above this concentration multimers of DNA molecules could only be captured. These multimers likely form by enhanced chain-chain association screenable by increasing NaCl concentration. Highlight curves of this dataset are shown in **Fig.2** of the main text.

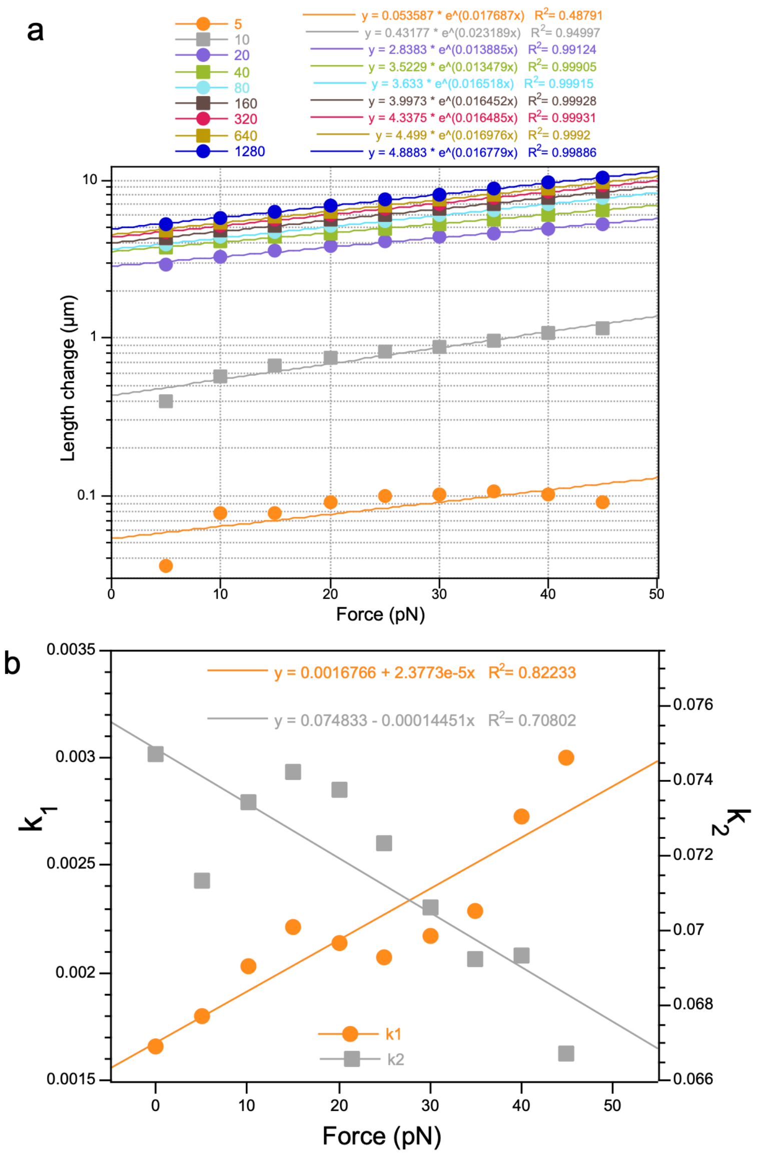


**Figure S2. a)** Length increment of dsDNA as a function of force at different TMPyP concentrations, in the presence of 0.01 M NaCl and at a pulling rate of 0.2 µm/s. The length change was calculated by subtracting the control (0 TMPyP) DNA length from the TMPyP-treated length measured at the given force. Data were fitted with single-exponential functions indicated in the legend. The figure shows the full dataset, part of which is shown in **Fig.4b** of the main text. **b)** Effect of force on the apparent rate constants ($k_1$ and $k_2$) of the double-exponential fits to the Length change versus TMPyP concentration curves shown in **Fig.4c** of the main text. The data were fitted with linear functions.

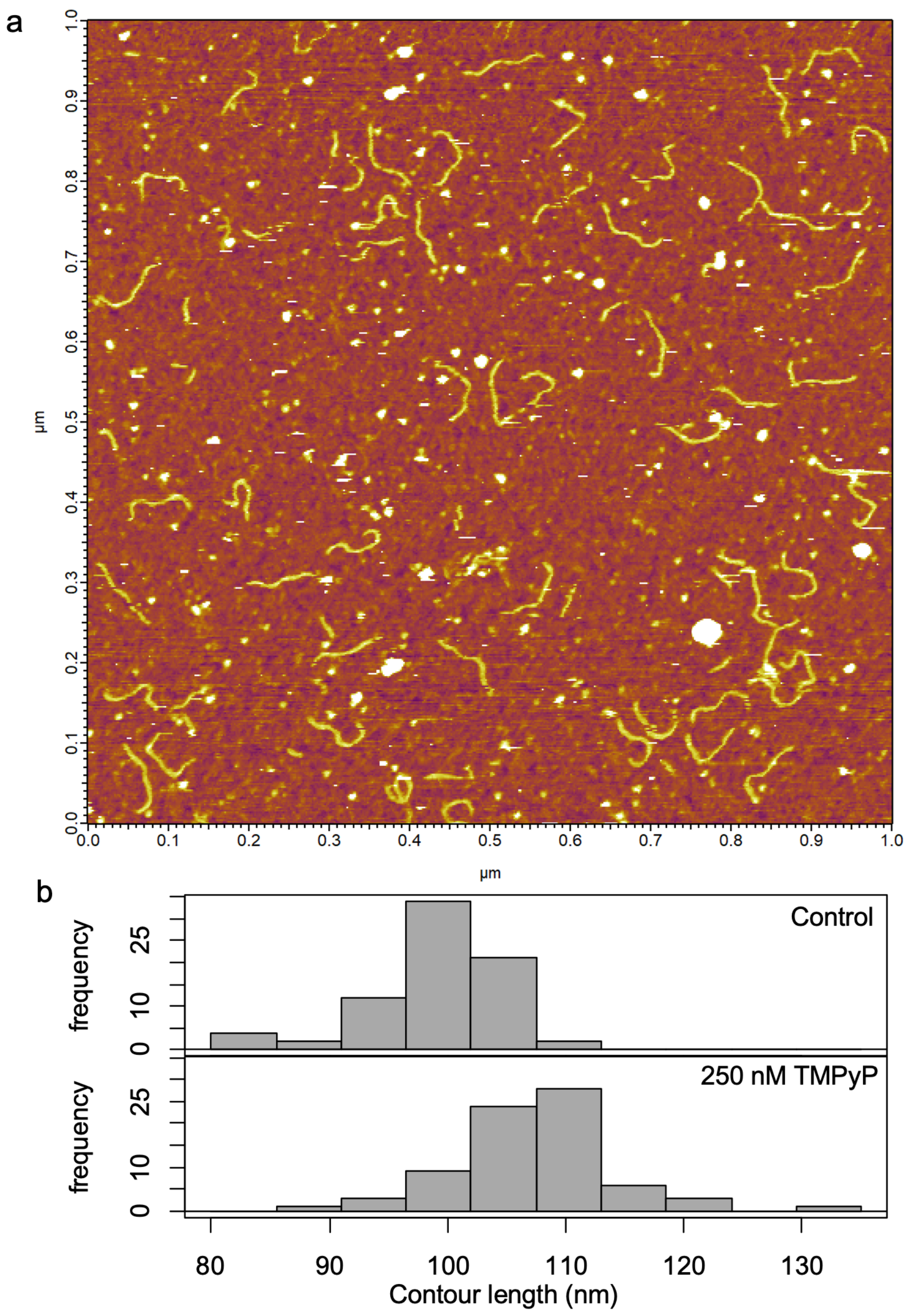


**Figure S3.** Effect of TMPyP on the contour length of conformationally relaxed dsDNA. **a)** AFM image of 300-bp-long dsDNA molecules equilibrated to poly-l-lysine-coated mica surface and treated with 250 nM TMPyP. **b)** Distribution of the contour length of control (top) and TMPyP-treated (bottom) dsDNA molecules. The mean contour lengths of the control and TMPyP-treated DNA molecules were 99.38 nm (±5.52 nm S.D., n=75) and 107.67 nm (±6.65 nm S.D., n=75), respectively. The mean contour lengths are significantly different (p-value 6.734e-14).

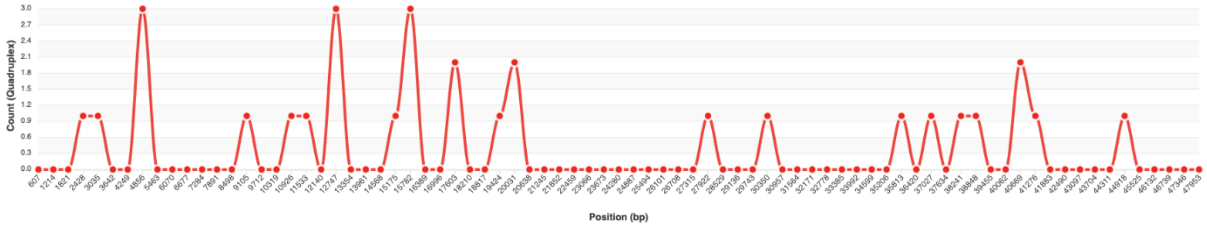


**Figure S4.** Distribution of G-quadruplex sequences in λ-phage DNA. The analysis was carried out by using the G4 hunter algorithm of the DNA analyser program package (https://bioinformatics.ibp.cz/#/analyse/quadruplex).

**Supplementary Video.** A single λ-phage DNA molecule stretched and relaxed in consecutive mechanical cycles with progressively changing maximal stretch length. 320 nM TMPyP, 1 M NaCl, 20 μm/s pulling rate. The video was generated by using a user-developed Python script based on the experimentally obtained data points. In the video the data points appear at the realistic time scale.
